# A quasi-integral controller for adaptation of genetic modules to variable ribosome demand

**DOI:** 10.1101/336271

**Authors:** Hsin-Ho Huang, Yili Qian, Domitilla Del Vecchio

## Abstract

The behavior of genetic circuits is often poorly predictable. A gene’s expression level is not only determined by the intended regulators, but also largely dictated by changes in ribosome availability imparted by activation or repression of other genes. To address this problem, we design a quasi-integral biomolecular feedback controller that enables the expression level of any gene of interest (GOI) to adapt to changes in available ribosomes. The feedback is implemented through a synthetic small RNA (sRNA) that silences the GOI’s mRNA, and uses orthogonal extracytoplasmic function (ECF) sigma factor to sense the GOI’s translation and to actuate sRNA transcription. Without the controller, the expression level of the GOI is reduced by 50% when a resource competitor is activated. With the controller, by contrast, gene expression level is practically unaffected by the competitor. This feedback controller allows adaptation of genetic modules to variable ribosome demand and thus aids modular construction of complicated circuits.

## Introduction

The ability to create complicated systems from composition of functional modules is critical to the progress of synthetic biology, yet it has been a longstanding challenge in the field [1–3]. Although progress has been made toward this goal [2–6], context-dependent behavior of genetic circuits is still a major hurdle to modular design [1, 3]. This leads to a combinatorial design problem where one has to re-design all circuit’s components every time a new component is added to a system. While many factors contribute to context-dependence of genetic modules, sharing limited gene expression resources has appeared as a major player into this problem [7–13].

In bacteria, the rate of gene expression is mainly limited by the availability of ribosomes [7, 8, 13]. In particular, activation of one transcriptional (TX) device, that is, a system where input transcriptional regulators affect expression level of one output protein, reduces availability of ribosomes to other, otherwise unconnected, TX devices, affecting their gene expression levels by up to 60% [7, 11]. These unintended non-regulatory interactions among TX devices can significantly alter the emergent behavior of genetic circuits. For example, the dose-response curve of a genetic activation cascade can be biphasic or even monotonically decreasing, instead of being monotonically increasing as expected from the composition of its TX devices [9]. A specific cascade’s response can be obtained by tuning the strength of unintended interactions that arise due to resource competition, which can be effectively accomplished by tuning ribosome binding site (RBS) strengths [9]. While this approach can recover the intended behavior of a circuit once in its final context, it is plagued by the problem that the circuit’s behavior may still be poorly robust to variations in ribosome availability due to changes in the circuit’s context. Therefore, there is a general need to find solutions that make the expression level of a TX device adapt to changes in ribosome availability while keeping the input and output connectivity of the device unchanged. This would enable seamless and scalable composition of TX devices whose input/output (i/o) behaviors are more robust to context.

In traditional engineering systems, feedback control has played a critical role into making circuit’s components modular, i.e., into maintaining a desired i/o response despite disturbances. This enabled predictable and reliable composition of larger systems from subsystems [14]. Recently, the concept of feedback control has been employed in synthetic genetic circuits to increase robustness of gene expression to fluctuations in available resources. In particular, Hamadeh et al. carried out theoretical analysis to compare the performance limits of different feedback architectures to mitigate effects of resource competition at various levels of gene expression [15]. Specifically, they considered two main classes of feedback controllers: transcriptional (TX) and post-transcriptional (post-TX). In a TX controller, negative feedback decreases the transcription rate of the GOI, while in a post-TX controller the feedback inhibits translation rate. In the case where the major resource competed for is ribosomes, a post-TX controller is theoretically sufficient for robustifying gene expression in the face of resource fluctuations. Shopera et al. [16] designed a TX controller by co-designing the regulatory input and output protein of a TX device such that the input can be sequestered by the anti-activator output protein, creating a negative feedback loop. As a result, the expression level of the TX device’s gene is more robust to fluctuations in availability of ribosomes. Although a promising proof of concept, the requirement to co-design the TX device’s input and output to engineer the feedback prevents generalization and scalability of this solution beyond the specific circuit’s instance considered.

In this paper, we design a post-TX feedback controller in which alteration of a TX device’s intended regulatory input is not necessary to engineer adaptation to changes in ribosome availability. Specifically, we designed and implemented a biomolecular feedback controller mediated through synthetic small RNA (sRNA) [17] to enable adaptation of the TX device’s output protein concentration to changes in ribosome availability. The key innovation of our solution is a quasi-integral control (QIC) strategy, which can approximate ideal integral control when all reactions constituting the controller are sufficiently faster than molecule decay [18]. Integral controllers are often responsible for homeostasis and perfect adaptation in natural systems [19, 20] and are ubiquitous in traditional engineering systems to ensure robustness to disturbances [21]. While genetic circuit motifs implementing integral control have been considered [22], they may be difficult to realize due to decay (i.e., dilution and degradation) of the biomolecules implementing the control reactions, which leads to integrator “leakiness” [18, 23, 24]. QIC manages this physical constraint by implementing all control reactions with sufficiently large reaction rates that are tunable through a “gain” parameter, mitigating the effects of integration leakiness. We demonstrate that when the sRNA feedback controller is absent, the output of the TX device is reduced by up to 50% when a resource competitor is activated. In contrast, when the sRNA feedback controller is present and configured to operate with high gain, the regulated TX device’s output becomes essentially unaffected by competitor activation, showing perfect adaptation.

## Results

### Enforcing modularity through embedded feedback control

A genetic circuit is commonly viewed as the composition of TX devices, which are systems that take transcriptional regulators (TX regulators) as inputs and produce proteins (possibly also TX regulators) as outputs. Modularity, the property according to which the i/o behavior of a system does not change with its context, is required for bottom-up design of synthetic genetic circuits [25]. However, the salient properties of a TX device often depend on context, which includes both direct connectivity to and mere presence of other TX devices [1, 3]. While problems of loads due to direct connectivity have been addressed in earlier works [4, 5], the problem of context-dependence due to resource sharing remains largely open [7, 9–12]. In particular, with reference to Figure 1A, the demand for ribosomes imparted by one TX device (TX device 2) in response to its regulatory input (*u*_2_) leads to a change *d* in the concentration of available ribosomes. This, in turn, affects the output (*y*_1_) of a different TX device (TX device 1), despite its intended regulatory input (*u*_1_) is unchanged (Figure 1B). As a result, the two TX devices become indirectly connected, breaking modularity and confounding circuit design.

**Figure 1:**
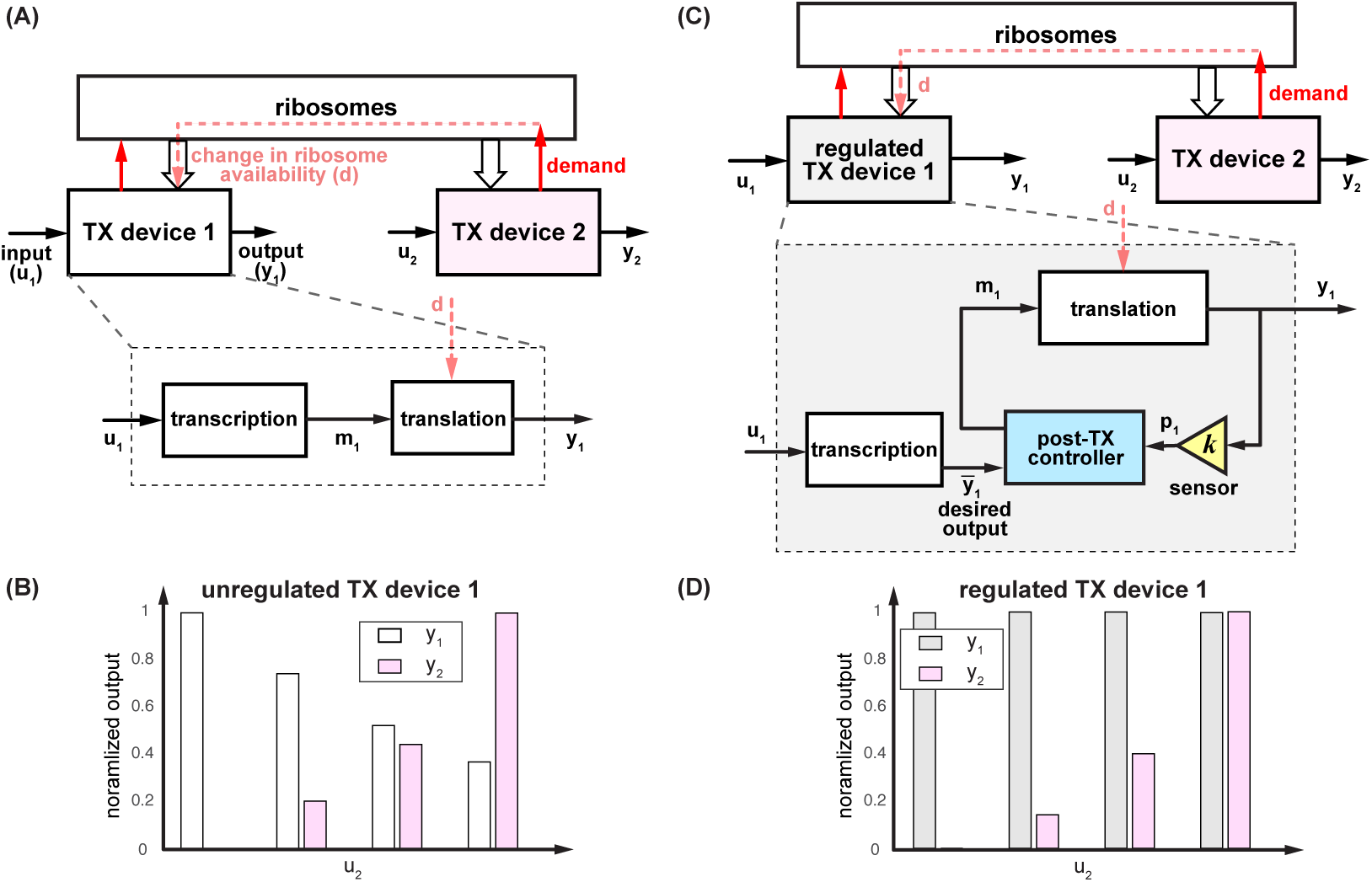
Modularity in resource-limited genetic circuits. (A) Each TX device *i* (*i* = 1, 2) takes a TX regulator as input (*u_i_*) and produces a protein as output (*y_i_*). When multiple TX devices share a pool of limited ribosomes, the demand from one device causes a change in ribosome availability, which consequently affects translation in other devices, creating unintended interactions among TX devices. (B) For the two-device circuit depicted in panel A, output from TX device 1 (*y*_1_) becomes coupled to the input to TX device 2 (*u*_2_), resulting in a circuit that lacks modularity. (C)-(D) By embedding a post-TX controller in a TX device, its output can adapt to changes in ribosome availability (panel D), eliminating the unintended interaction between devices due to ribosome competition, thus rescuing modularity. The post-TX controller does not alter a TX device’s intended regulatory input nor the output and can therefore enable seamless composition of regulated TX devices.

To restore modularity of TX device 1, we engineer a post-TX feedback controller embedded in this device to make its i/o behavior independent of TX device 2 (Figure 1C). This feedback controller senses the output *y*_1_ of TX device 1 and compensates for the effect of a drop or an increase in ribosome concentration (*d*) by appropriately adjusting the mRNA level *m*_1_. As a result, under a fixed input *u*_1_, the output *y*_1_ of TX device 1 adapts to changes in ribosome demand by TX device 2 (Figure 1D). Since this feedback design is post-TX, it is also orthogonal to the intended regulatory input *u*_1_ of TX device 1, enabling composition of multiple TX devices through desired TX regulatory interactions.

The problem of making the output of TX device 1 independent of TX device 2’s increased demand for ribosomes can be solved by regarding the variation in ribosome availability *d* to TX device 1 as a disturbance and solving the control theoretic problem of disturbance rejection [21]. Rejection of (i.e., perfect adaptation to) a disturbance can be accomplished by integral feedback control. An integral controller computes the difference between the desired output 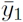 and the measured output *p*_1_ = *ky*_1_ as the error 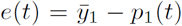. A memory element *z* in the controller accumulates (i.e., integrates) this error over time: 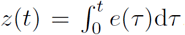. The memory of the error *z* is then used to determine the amount of actuation (i.e., mRNA *m*_1_) applied to the process to be regulated (i.e., translation). If the feedback interconnection of the process to be regulated and the integral controller is stable, then at steady state, we have that d*z/*d*t* = 0, which leads to *e* = 0 regardless of the disturbance input *d*. Due to this property, integral control is applied ubiquitously to engineering systems (e.g., vehicle cruise control systems [21]) and identified in natural networks (e.g., bacterial chemotaxis [19] and calcium homeostasis [20]). Therefore, we aim to design a post-TX controller (Figure 1C) that can achieve integral-control-like behavior to render the output of TX device 1 completely independent of TX device 2.

### Design of a sRNA-mediated post-TX controller

In order to evaluate the ability of the controller to make the output of TX device 1 adapt to changing ribosome demand by TX device 2, we assembled a test-bed genetic circuit with two TX devices shown in Figure 2A. Specifically, since TX device 2 needs to apply a variable demand for ribosomes, we made TX device 2 externally inducible and made its output be a red fluorescent protein (RFP) to be able to assess ribosome demand. We embedded a post-TX controller enabled by sRNA silencing in TX device 1. For this device, we chose a constitutive promoter since our design needs to demonstrate that the TX device’s output (GFP level) stays unchanged when the device’s input is kept constant, despite a change in ribosome availability. This is a model system for studying competition for ribosomes as employed in earlier works [7, 11, 16, 26]. Specifically, referring to Figure 2A, a pLacIQ promoter is used to constitutively express LuxR. In the presence of LuxR’s effector *N*-hexanoyl-l-homoserine lactone (AHL), AHL-bound LuxR (*u*_2_) is formed to activate TX device 2 to produce RFP (*y*_2_). These two genes constitute a *resource competitor* that demands more ribosomes when AHL concentration increases, leading to a decrease in the amount of available ribosomes to translate mRNA in TX device 1.

**Figure 2:**
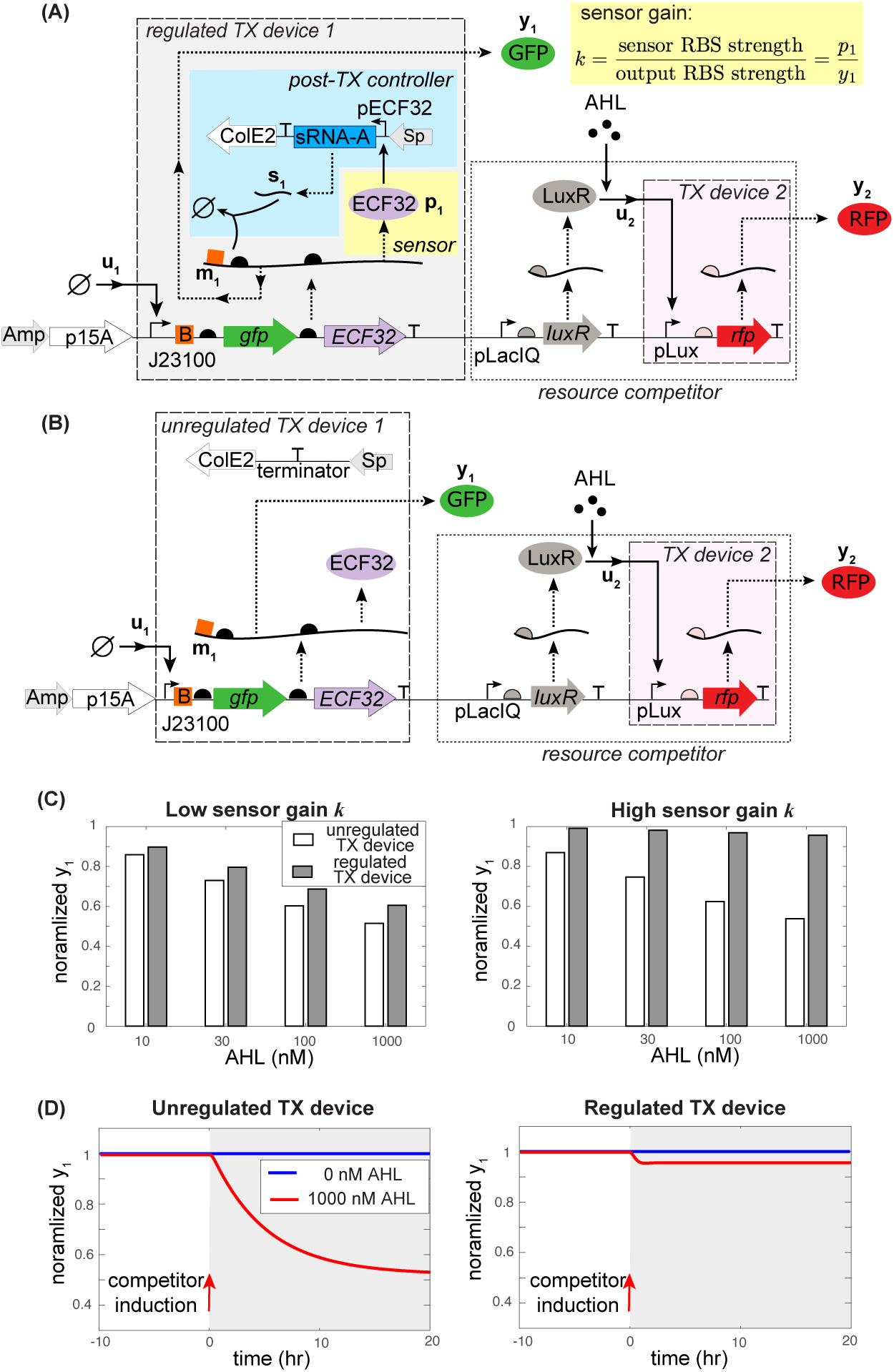
sRNA-mediated quasi-integral post-TX controller. (A) A regulated TX device 1 with inducible TX device 2 functioning as a resource competitor. Upwards arrow with tip rightwards indicate promoters, semicircles represent RBSs, orange box with letter B stands for sRNA-A targeting sequence, and “T” symbols represent terminators. The feedback controller consists of ECF32 as a sensor for GFP concentration, which is used to actuate transcription of sRNA-A. sRNA-A antisenses its targeting sequence on the mRNA for degradation of both RNA molecules. (B) Unregulated TX device 1 with inducible TX device 2 functioning as a resource competitor. The pECF32 promoter and sRNA-A in panel A are removed, and therefore, there is no feedback controller in this TX device. (C) Simulations of equations (1)-(3) for steady state behavior of the circuits in panel A-B with different competitor induction levels and sensor gains (*k*). (D) Simulation of the temporal response of GFP output from the unregulated TX device (circuit in panel B) and the regulated TX device (circuit in panel A). The resource competitor is induced at *t* = 0 with AHL concentrations indicated in the legend. Simulation parameters are listed in SI Section B6.

TX device 1 embeds the sRNA-mediated post-TX controller. The controller consists of four key biological parts: sRNA-A and its targeting sequence [17], extracytoplasmic function (ECF) sigma factor 32 1122 (referred to as ECF32 here after), and its cognate promoter pECF32 [27]. The choice of ECF32 and pECF32 is based on their minimal impact on host cell growth and their dose response’s wide dynamic range (see SI Section A2). In particular, *ECF32* gene is introduced downstream of the output *gfp* gene to form a bi-cistronic operon. This operon is driven by the strong constitutive BioBrick promoter BBa J23100 to co-transcribe *gfp* and *ECF32* genes. Since translation of ECF32 and GFP share the same pool of ribosomes, concentration of ECF32 translated is a proxy of GFP concentration. Therefore, with reference to Figure 1C, ECF32 can be regarded as a sensor that senses GFP concentration to actuates sRNA transcription. In fact, by a simple ribosome sharing model [9], the sensor (i.e., ECF32) concentration is proportional to the output (i.e., GFP) concentration by a sensor gain *k*, which can be quantified by the ratio between the ECF32’s and the GFP’s RBS strength (see Figure 2A and SI Section B1). ECF32 transcribes sRNA-A through its cognate promoter pECF32 on a separate plasmid. The mRNA co-transcript of GFP and ECF32 bears the sRNA-A’s targeting sequence immediately upstream of the *gfp* gene’s RBS. The sRNA can complementarily pair with its target mRNA, forming an inert RNA complex that is rapidly cleaved by RNase E, leading to coupled degradation of both mRNA and sRNA [28–32]. When the resource competitor is activated, the amount of ribosome available to translate GFP and ECF32 decreases, causing a reduction in their concentrations. Reduction in ECF32 concentration reduces sRNA transcription, which consequently increases the amount of GFP/ECF32 mRNA co-transcripts to recover their translation activity, closing the feedback loop.

To experimentally evaluate the benefit of this sRNA-mediated post-TX controller, we constructed another circuit with unregulated TX device 1 and the resource competitor (Figure 1B). In particular, the only difference is the absence of pECF32 promoter and sRNA-A message in the unregulated device. As a result, the mRNA level in the unregulated TX device 1 (*m*_1_) is not responsive to output *y*_1_, breaking the feedback loop. This circuit allows us to compare the robustness of the regulated and the unregulated TX device 1 subject to the same level of competitor induction to quantify the performance of this post-TX controller.

### Mathematical analysis of the regulated TX device’s adaptation property

With reference to Figure 2A, the biomolecular reactions involved in the regulated TX device 1 can be described by the following mass-action kinetic model:

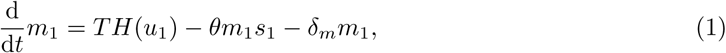

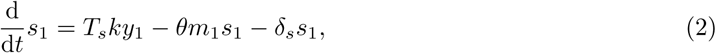

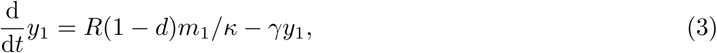
where *m*_1_, *s*_1_ and *y*_1_ stand for the concentrations of the GFP/ECF32 mRNA co-transcript, sRNA, and GFP protein, respectively. Function *H*(*u*_1_) describes transcriptional regulation by input *u*_1_ and is a constant when the TX device’s promoter is constitutive. Parameter *k* is the sensor gain defined in Figure 2A; *T* and *T_s_* are the transcription rate constants of the mRNA and the sRNA, respectively, which are proportional to their DNA copy numbers and promoter strengths; *θ* is the degradation rate constant of the mRNA-sRNA complexes; *δ_m_* and *δ_s_* are the respective decay (i.e., degradation and dilution) rate constants of uncoupled mRNA and sRNA; *γ* is the dilution rate constant; *R* is the maximum translation rate constant proportional to the total amount of ribosomes; *κ* is the RBS strength of GFP and 0 ≤ *d* < 1 models a change in the availability of ribosomes. The models of RNA transcription, decay and RNA-RNA interaction in equations (1)-(2) are standard and can be found in [28, 33–35]. Nevertheless, we derived them from chemical reactions in SI Section B1.

If the uncoupled RNAs do not decay (i.e., *δ_m_* = *δ_s_* = 0), this sRNA-mediated feedback system is an antithetic integral controller (AIC) [22], which performs integral control. Specifically, the integral action in the RNA controller dynamics (1)-(2) appears through the memory variable *z* := *m*_1_ − *s*_1_, whose dynamics follow

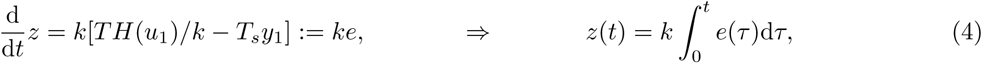
where the error *e*(*t*) is specified by the difference between the desired output 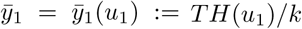, which is a function of the regulatory input *u*_1_ only, and the scaled output *T_s_y*_1_:

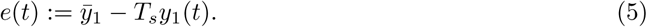

Suppose that the ODE model (1)-(3) is stable, then the time derivative of *z* in (4) reaches 0 at steady state, resulting in the steady state output to be 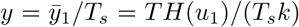, which is independent of the disturbance *d*. This implies that expression of the regulated TX device 1 (*y*_1_) is independent of (i.e., adapts perfectly to) the disturbance *d* and only depends on its own regulatory input *u*_1_.

However, due to (i) degradation by RNase and (ii) dilution due to fast growth of bacteria, decay of uncoupled mRNA and sRNA are unavoidable in practice [28] (*δ_m_,δ_s_* > 0). As a consequence, the memory variable dynamics become

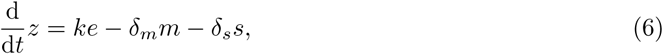
which, in contrast to equation (4), does not contain an integral of the error. In fact, the memory *z* gradually fades away due to decay of RNA species. This disruption of the key integral control structure can result in potentially large adaptation error, making it practically impossible for the output *y*_1_ to adapt perfectly to a variation *d* in ribosome availability.

Nevertheless, perfect adaptation can be asymptotically achieved by this sRNA-mediated controller by rational choice of circuit parameters. In fact, to overcome the undesirable effect of fading memory, one can engineer the accumulation of the error to take place at much faster rates than the rates at which uncoupled mRNA and sRNA decay. This allows the memory signal to be amplified, making its decay negligible, and thus leading to practically no adaptation error. This strategy is called quasi-integral control (QIC) and a more in-depth mathematical analysis of its properties and stability can be found in [18] and SI Section B2-3. From equation (6), it follows that in the sRNA-mediated controller, the rate at which error *e* is accumulated is determined by the gain parameter *k*. Therefore, in order to enforce QIC in the sRNA-mediated controller, we can increase the sensor gain *k* by increasing the RBS strength of ECF32 (see Figure 2A and detailed analysis in SI Sections B2).

These model predictions are confirmed through simulations in Figure 2C. In particular, we find that while the output of the unregulated TX device 1 changes significantly when ribosome availability drops, the output of the regulated TX device 1 is less sensitive to variations in ribosome availability. This implies that sRNA-mediated feedback can make gene expression in a regulated TX device adapt to variations in ribosome availability, and as a consequence, the output of TX device 1 becomes completely independent of TX device 2. Simulations in Figure 2C also confirm that the adaptation error of a regulated TX device can be effectively quenched by increasing the sensor gain *k*, leading to essentially perfect adaptation to changes in ribosome availability and therefore to an entirely modular TX device. In Figure 2D, we simulated the temporal response of a QIC subject to a step induction of the resource competitor. We find that due to the fast reactions in the QIC, adaptation occurs at a timescale that is much faster than gene expression.

### Experimental results: Regulated TX module adapts to changes in ribosome demand

Before testing the circuits in Figure 2A-B, we first verified that sRNA silencing mechanism can inhibit GFP expression as expected. To this end, we performed a GFP silencing experiment using the circuit in Figure 3A (see plasmid maps in SI Figure S2). Specifically, the mRNA of *gfp* gene to be silenced has sRNA-A’s targeting sequence (orange B box) located immediately upstream of the gene’s RBS and it is transcribed by a strong constitutive promoter BBa J23100 from the BioBrick Registry. sRNA-A is transcribed by pECF32 promoter in the presence of ECF32, whose expression is regulated by TetR repressor and its effector aTc. We confirmed that increasing the concentration of aTc decreases GFP expression, resulting in GFP expression to be reduced by about 6-fold. In contrast, when pECF32 promoter and sRNA-A message were removed, we observed high levels of GFP expression regardless of aTc concentration (Figure 3B). These results confirm that the sRNA-A can silence GFP expression as desired.

**Figure 3:**
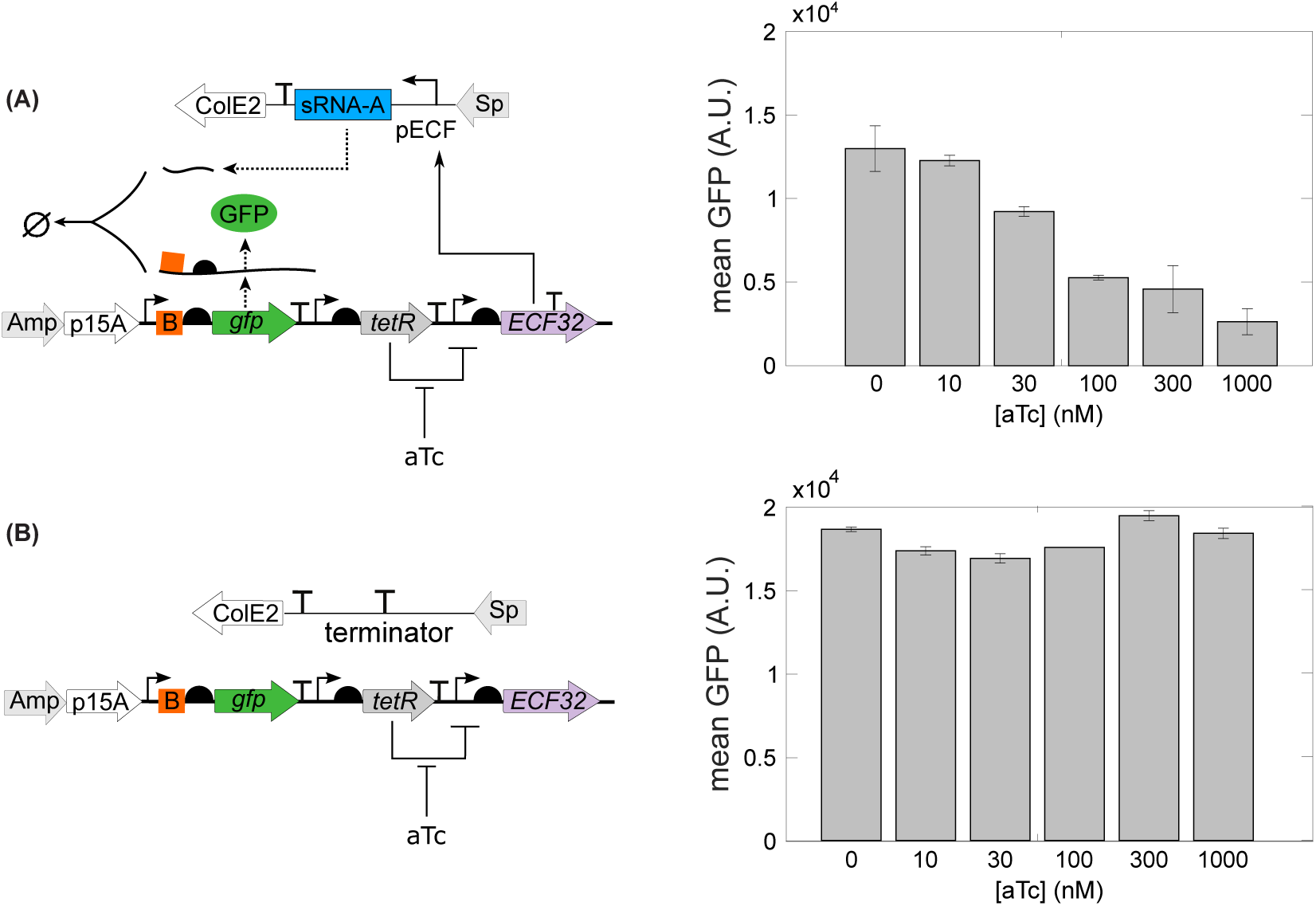
Genetic circuits to test the sRNA-A silencing and the ECF32 actuation mechanisms. (A) GFP expression is regulated by the sRNA-A silencing and the ECF32 actuation mechanisms. When TetR-regulated ECF32 is induced by TetR’s effector aTc, ECF32 binds to its cognate promoter pECF32 to transcribe sRNA-A. sRNA-A then antisenses to its targeting sequence (orange B box) on GFP’s mRNA to silence GFP expression. The mechanisms to be tested were confirmed by a 6-fold GFP reduction when adding aTc. (B) When pECF32 promoter and sRNA-A message were removed, GFP expression was not affected by aTc concentration. This further confirmed the mechanisms to be tested. Data in both panels were obtained with microplate photometer. The error bars are standard error of mean from three replicates.

Having verified the sRNA silencing and the ECF32 actuation mechanisms, we constructed the circuit in Figure 2A(B) that consists of a resource competitor and a regulated (unregulated) TX device. The plasmid maps and DNA sequences can be found in SI Figure S3 and SI Section A3, respectively. Given that from the mathematical analysis and the simulation in Figure 2C, the RBS strength of ECF32 is a key parameter responsible for the sensor gain and the adaptation error of this sRNA-mediated QIC, we created a library of genetic constructs with different RBS strengths for ECF32 (see SI Table S3). We first employed a weak RBS for ECF32 with a translation initiation rate (TIR) of 471 calculated by the RBS calculator 2.0 [36]. As shown in Figure 4A, while GFP output from the unregulated TX device was reduced by about 50% when AHL was increased to 1000 nM, GFP output from the regulated TX device is only reduced by 20%. To completely eliminate any dependence of the output of regulated TX device 1 on the operation of TX device 2, we followed the model prediction in Figure 2C to increase the sensor gain by employing a strong RBS with a TIR of 6471 for ECF32 in both the regulated and the unregulated TX device. We observed that while GFP output from the unregulated TX device was reduced by up to 50% when 1000 nM of AHL was applied to induce the competitor, GFP expression from the regulated TX device was essentially unaffected by AHL induction (Figure 4B). In Figure 4C-D, we recorded the temporal response of the unregulated TX device and the regulated TX device with high sensor gain. TX device 2 was activated at time 0 with 0 or 1000 nM AHL. The temporal responses are consistent with simulation results in Figure 2D, where the unregulated TX device’s output drops after AHL induction and does not recover without the controller. By contrast, with the controller, the output from the regulated TX device immediately recovers from the applied perturbation. These results demonstrate that the sRNA-mediated QIC is an effective approach to insulate the behavior of a TX device from its surrounding systems and to maintain modularity of genetic circuits.

**Figure 4:**
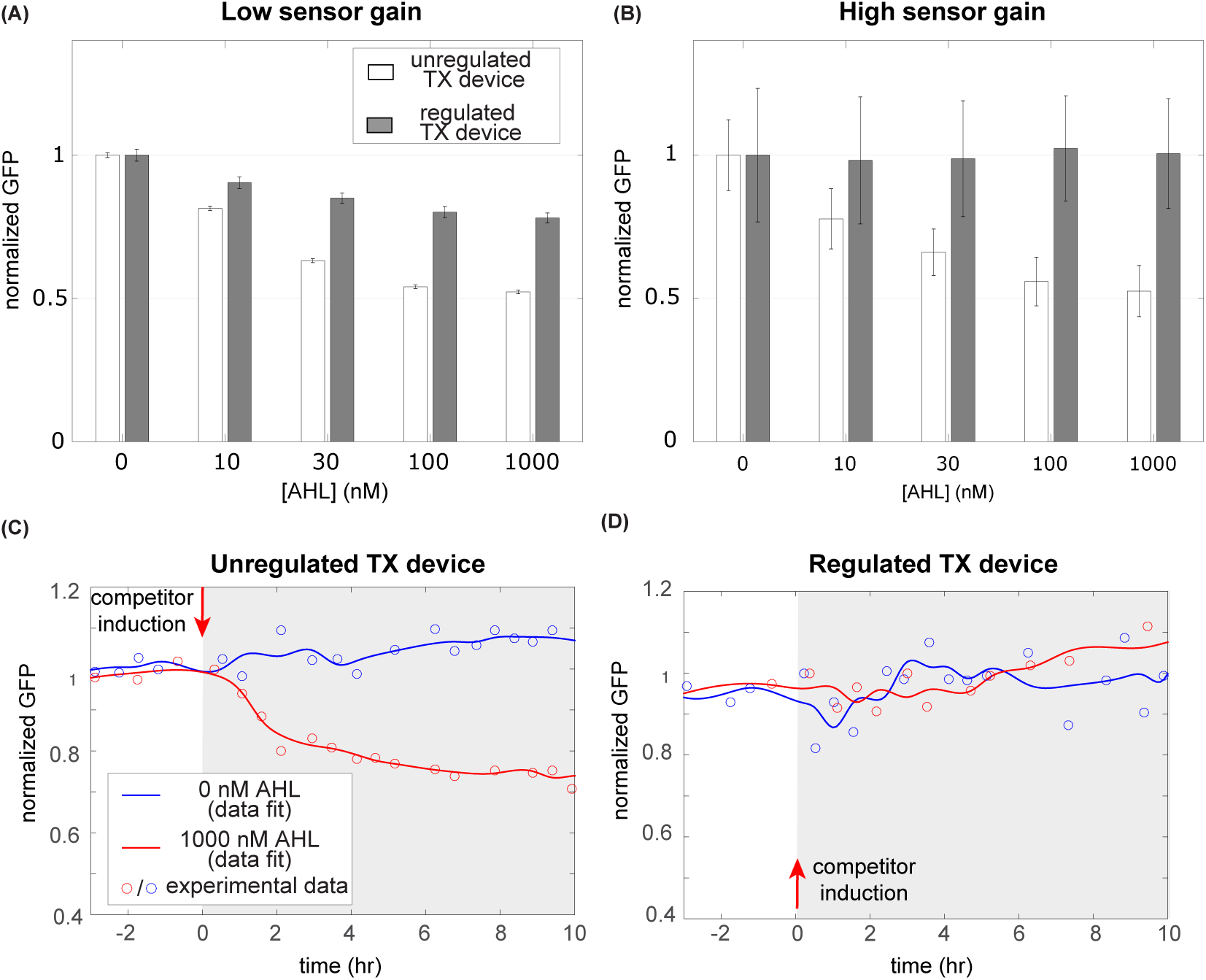
A regulated TX device with sRNA-mediated QIC can adapt perfectly to changes in ribosome availability. (A)-(B) Normalized steady state GFP expression from the regulated TX device (circuit diagram in Figure 2A) and the unregulated TX device (circuit diagram in Figure 2B), with different levels of AHL induction of the resource competitor. GFP expression levels were normalized to its mean values when the resource competitor is not induced (i.e., AHL=0 nM). The RBS of the ECF32 sensor has a TIR of 471 (corresponding to low sensor gain) in panel A and a TIR of 6471 (corresponding to high sensor gain) in panel B. Data were obtained with microplate photometer. The error bars are standard error of mean of GFP per OD from three replicates. See SI Figure S6 for unnormalized data. (C)-(D) Temporal response of GFP output from the unregulated TX device (circuit in Figure 2B) and the regulated TX device (circuit in Figure 2A). The resource competitor was induced at *t* = 0 with AHL concentrations indicated in the legend. GFP expressions were normalized to their respective levels at *t* = 0. Experimental data (shown by circles) were acquired from multiplex turbidostat and automatic flow cytometry in a single measurement. The fitting curves were obtained by a first order moving average filter and fitted by cubic spline interpolation. See SI Figure S7 for unnormalized data.

## Discussion

Modular design of genetic circuits relies on the assumption that the input/output behavior of TX devices is not affected by surrounding systems. This allows creation of increasingly sophisticated systems, whose behavior can be conveniently predicted by that of the composing devices characterized in “isolation”. While substantial progress has been made toward making the input/output behavior of a device independent of its context [3, 4, 6], a TX device’s response can still be severely affected by operation of other devices through resource sharing [7–13]. This impedes a modular approach to designing complicated systems, hampering progress in synthetic biology. In this paper, we have proposed a solution to this problem, which allows the output response of a TX device in bacteria to adapt to changes in availability of ribosomes imparted by other systems. Our solution is based on embedding a post-TX feedback controller within a TX device, which treats changes in ribosomes concentration as a disturbance. The controller reaches perfect adaptation by implementing a quasi-integral control (QIC) action [18] through sRNA-mediated mRNA silencing. In a QIC scheme, ideal integral action and hence perfect adaptation can be asymptotically reached as a control gain parameter is progressively increased. Here, we have demonstrated that this parameter can be easily tuned through the ECF sigma factor’s RBS strength, leading to a practical way to tune controller’s performance (Figure 4).

Chemical reactions with similar structure to sRNA-mediated silencing have been studied by others and have been proposed as a way to implement integral control under the assumption that molecule decay can be neglected [22]. While this assumption is reasonable in slow-growing cells and with controller species with negligible degradation, our experimental results suggest that this may not always be the case for uncoupled RNA species in *E. coli*. In fact, under the assumption of negligible decay, perfect adaptation should be achieved regardless of the feedback gain. By contrast, our controller reaches perfect adaptation only for sufficiently large controller gain (Figure 4A-B), suggesting QIC instead of a pure integral control structure. To approach perfect adaptation, we increased the feedback gain by increasing the RBS strength of the sensor protein (i.e., ECF32), which increases its concentration to promote sRNA transcription rate. We have characterized the promoter pECF32 and found that it has a wide dynamic range (see SI Section A2). This is an important aspect of the design as a narrow dynamic range may lead to saturate a ECF promoter when its cognate ECF sigma factor’s RBS strength is increased, breaking the feedback loop (see SI Section B4).

In contrast to other efforts aimed to regulate gene expression by modulating its TX rate [16, 37] our control design is post-TX and hence completely orthogonal to TX regulation. It thus enables seamless i/o composition of TX devices, consistent with a modular design concept. Compared to other sRNA-based circuits [38], our controller’s constituent biomolecular parts, specifically the library of sRNA-As and ECF sigma factors, are highly expandable and interchangeable [17, 27], therefore this feedback design is a generalizable and scalable solution to engineer modular genetic circuits with increased size and complexity. While the approach of this paper can be regarded as a decentralized control approach, wherein feedback controllers are built in TX devices, an orthogonal line of research has focused on a centralized control approach. With reference to Figure 1A, in this line of research, solutions globally manipulate resource pool size and/or allocation. For instance, orthogonal ribosomes have been proposed to decouple resource demand by the host from that by synthetic circuits [39]. Yet, unintended interactions arising from synthetic TX devices remains a problem, and there currently does not exist a sufficiently large library of orthogonal ribosomes to assign an independent resource to each TX device. In [40], Venturelli et al. engineered a sequence-specific ribonuclease to cleave certain host cell transcripts, thus diverting resources to synthetic proteins to increase productivity of synthetic metabolic pathways. Conversely, Ceroni et al. constructed a dCas9-based feedback controller that represses the synthetic circuit when its resource demand is high, thus reallocating resources to maintain constant host cell growth [41]. In [26], Darlington et al. partitioned the ribosome pool into synthetic circuit-specific and host-specific ribosomes using orthogonal rRNAs, and implemented a feedback controller to increase the portion of synthetic circuit-specific ribosomes when more are demanded. However, depending on the genetic circuit’s topology, more abundant ribosomes can actually be deleterious and can increase the relative strength of unintended interactions due to resource sharing (SI Section B5). Therefore, what solution to use may largely depend on the application, with a decentralized solution being especially promising in applications involving signal transduction and logic computation, where accuracy and precision, rather than high yield, are the main design considerations. For future large integrated genetic circuits that encompass both metabolic pathways and signal transmission, a combination of both centralized and decentralized solutions may be optimal.

Homeostasis and adaptation has long been argued as a key property for survival of living organisms in highly uncertain and variable environments [42]. Several sRNA feedback motifs similar to ours have been identified in natural systems to regulate key physiological pathways. For example, an alternative sigma factor *σ^E^* controls extracytoplasmic stress response in *E. coli*. *σ^E^* is required to produce sRNA RybB, while being repressed by it [43]. Negative feedback regulation through sRNA is also involved in iron homeostasis in *E. coli* [44–46], in quorum-sensing response [47, 48], and in sugar metabolism [49, 50]. Given our results, instances of such a feedback motif in natural biological pathways may hint to a previously under-appreciated ability of such pathways to withstand perturbations.

## Methods

### Strain and growth medium

*Escherichia coli* DIAL strain JTK-160J [51] was used in all the constructions and experiments in this work. The growth medium is M9610 medium supplemented with 0.4 % glucose, 0.2% casamino acids, and 1 mM thiamine hydrochloride. Appropriate antibiotics were added according to the selection marker of a genetic circuit. Final concentration of ampicillin or spectinomycin is 100 µg mL^−1^. M9610 minimal medium was modified from M9 minimal medium to make phosphate buffer pH 6 and its buffer capacity 10-fold of the one of M9 minimal medium according to the Henderson-Hasselbalch Equation. The recipe of M9610 minimal medium is Na_2_HPO_4_ · 7H_2_O 33.36 mM, KH_2_PO_4_ 220.40 mM, NaCl 8.56 mM, and NH_4_Cl 18.69mM. The pH 6 and 30◦C growth condition has been shown to lower degradation rate of LuxR’s effector AHL [52].

### Genetic circuit construction

The genetic circuit construction was based on Gibson assembly [53]. DNA fragments to be assembled were amplified by PCR using Phusion High-Fidelity PCR Master Mix with GC Buffer (NEB, M0532S), purified in gel electrophoresis with Zymoclean Gel DNA Recovery Kit (Zymo Research, D4002), quantified with the nanophotometer (Implen, P330), and assembled with Gibson assembly protocol using NEBuilder HiFi DNA Assembly Master Mix (NEB, E2621S). Assembled DNA was transformed into competent cells prepared by the CCMB80 buffer (TekNova, C3132). Plasmid DNA was prepared by the plasmid miniprep-classic kit (Zymo Research, D4015). DNA sequencing used Quintarabio DNA basic sequencing service. The pVRa32 1122 and pVRb32 1122 plasmids for cloning ECF32 and its cognate promoter pECF32 were purchased from Addgene (ID 49689 and 49722, respectively). sRNA-A was synthesized in a gBlock. Primers and gBlocks were obtained from Integrated DNA Technologies. The list of primers and constructs is in SI Section A1.

### Microplate photometer

Pre-culture was prepared by inoculating a −80◦C glycerol stock in 650 µL growth medium per well in a 24-well plate (Falcon, 351147) and grew at 30◦C, 250 rpm in a horizontal orbiting shaker for 7 hours. Pre-culture was diluted to an initial OD at 600 nm of 0.001 in 200 µL growth medium per well in a 96-well plate (Falcon, 351172). The plate was incubated at 30◦C and was analyzed with OD, GFP, and RFP intensity in a Synergy MX (Biotek, Winooski, VT) microplate reader.

### Multiplex turbidostat and automatic flow cytometry

We devised a home-built system of multiplex turbidostat and automatic flow cytometry to grow bacterial cells in a continuous culture at maximal growth rate and online monitor single-cell fluorescence with flow cytometry automatically [54]. The multiplex bioreactor was based on Flexostat [55] and can run up to 16 tubes simultaneously. Pre-culture, as described in the microplate photometer experiment, was diluted to 0.1 of OD at 600 nm in a 1 mL growth medium as an inoculum. The inoculum was inoculated in a tube of the multiplex bioreactor to reach the working volume as 15 mL per tube. The multiplex bioreactor was accommodated in an incubator (VWR, Forced Air General Incubator, 414005-120) to control temperature at 30 ◦C. The OD set point of turbidostat is 0.1 at 650 nm. Online monitoring of single-cell fluorescence was achieved by the automatic flow cytometry. The flow cytometer Accuri C6 (BD Biosciences) was connected to a home-built autosampler modified from the MSP FlowCytoPrep 5000 model. The autosampler automatically transferred bacterial cells from a tube of the multiplex bioreactor to the SIP (sample injection port) of Accuri C6 and remotely controlled Accuri C6 to analyze the sample. Accuri C6 detected GFP and red fluorescent protein (RFP) emission with the band-pass filter 525/50 and 610/25, respectively. The data-collecting threshold is 10000 on FSC-H. The data was first gated in a FSC-A and SSC-A with a threshold of 10000 on FSC-A channel. Singlet cells were then gated in a FSC-A versus FSC-H plot and the collected events of singlets are about 226000 counts in average. Fluorescence data of singlets were analyzed with arithmetic mean. The current system’s sampling rate is about every 3.5 min per sample. Higher sampling rate is achieved by skipping the dilution step [54] because a low turbidostat OD set point as 0.1 allows the cytometer’s collecting rate in the linear range of analysis.

